# Tumor-infiltrating immune repertoires captured by single-cell barcoding in emulsion

**DOI:** 10.1101/134841

**Authors:** Adrian W. Briggs, Stephen J. Goldfless, Sonia Timberlake, Brian J. Belmont, Christopher R. Clouser, David Koppstein, Devin Sok, Jason Vander A. Heiden, Manu V. Tamminen, Steven H. Kleinstein, Dennis R. Burton, George M. Church, Francois Vigneault

## Abstract

Tumor-infiltrating lymphocytes (TILs) are critical to anti-cancer immune responses, but their diverse phenotypes and functions remain poorly understood and challenging to study. We therefore developed a single-cell barcoding technology for deep characterization of TILs without the need for cell-sorting or culture. Our emulsion-based method captures full-length, natively paired B-cell and T-cell receptor (BCR and TCR) sequences from lymphocytes among millions of input cells. We validated the method with 3 million B-cells from healthy human blood and 350,000 B-cells from an HIV elite controller, before processing 400,000 cells from an unsorted dissociated ovarian adenocarcinoma and recovering paired BCRs and TCRs from over 11,000 TILs. We then extended the barcoding method to detect DNA-labeled antibodies, allowing ultra-high throughput, simultaneous protein detection and RNA sequencing from single cells.

---

Sequencing of the immune repertoire has wide applications in basic immunology^,1^, autoimmunity^2^, infectious disease^3^ and oncology^4^. While many studies have investigated BCR and TCR diversity in circulating blood, there is growing interest in the immune receptors of TILs^5^, whose functions are highly relevant to cancer growth or regression yet variable and often uncharacterized. A critical step towards a better understanding of TILs will be recovery and functional characterization of their BCRs and TCRs, since this may allow the identification of new tumor-associated antigens. Tumor antigens are critically required to develop cancer vaccines^6^, understand the role of checkpoint inhibitors^7,8^, and advance chimeric antigen receptor T-cell (CAR-T^9^) therapy in solid tumors. However, despite decades of technical progress in immune sequencing^10–12^, no technology has been able to recover full-length, natively paired BCRs (heavy and light chains) and TCRs (alpha and beta chains) from a heterogeneous sample such as a tumor without *in vitro* culture or cell sorting, steps that restrict and bias the observed repertoire^13,14^. Several recent studies have demonstrated important technological advances toward this goal, but the reported methods fall short of the requirements often needed for immune repertoire analysis, particularly of complex heterogeneous material such as dissociated tumors. A recent study reported high throughput pairing of TCR alpha and beta receptor sequences based on repeated observations of TCR pairs across wells of a 96-well plate^15^. While powerful, since this is not a single cell approach, data recovery is limited to the variable TCR genes and only from clonally expanded TCR lineages. In a different approach, Dekosky et al.,^11^ demonstrated recovery of paired immunoglobulin heavy and light chain genes from single cells employing emulsion overlap PCR to link the paired amplicons. However, this method is limited to partial variable region sequence recovery, and due to its amplicon linking approach can only recover the BCR genes and no additional features of the cells. Outside immune sequencing, several studies have reported compelling progress in scale-up single cell whole transcriptome analysis^16–19^. In particular, the methods of Klein et al.^16^ (InDrop) and Macosko et al.^17^ (Dropseq) allow low-coverage transcriptome sequencing from several thousand cells. However, capture of full-length immune receptor genes presents a substantially different challenge to low-pass transcriptome tag counting. Firstly, immune sequencing requires capture of the full length variable region of BCR and TCR genes, which consist of several hundred base pairs of sequence extending all the way to the 5’ end of the transcripts. This differs substantially from the capture of short 3’ tags close to the poly-A tail as performed by existing emulsion single-cell RNAseq platforms. Secondly, meaningful immune repertoire analysis requires processing of hundreds of thousands to millions of cells^5^, especially if the input material such as dissociated tumor cells is unsorted, since only a minority of the input cells will be lymphocytes, representing an order of magnitude higher throughput requirement than has been demonstrated by single cell emulsion RNA sequencing methods to date.

In order to allow comprehensive analysis of natively paired BCRs and TCRs together with other genetic markers from complex heterogeneous samples we developed a microfluidic emulsion-based method for parallel isolation and DNA barcoding of millions of single cells (Figure 1). Up to a million cells per hour are isolated in individual ∼65 picoliter emulsion droplets, which are ∼15-75 times smaller than those used in recent single-cell transcriptome studies ^16,17^, therefore allowing correspondingly higher throughout. Within the droplets cells are lysed (Suppl. Fig. 1), target mRNA is reverse transcribed with target-specific primers and a two-step DNA barcoding process attaches both molecule-specific^20,21^ and droplet-specific barcodes to the cDNAs. After subsequent recovery and next generation sequencing, the dual barcoding strategy allows clustering of sequence reads into both their molecules and cells of origin. This allows extensive correction of errors and amplification biases^21^, clone counting at both the mRNA and cellular levels, heavy chain isotype determination, and importantly, recovery of full-length, natively paired V(D)J sequences of BCR and TCRs simultaneously at extremely high throughput.

**Figure 1.**
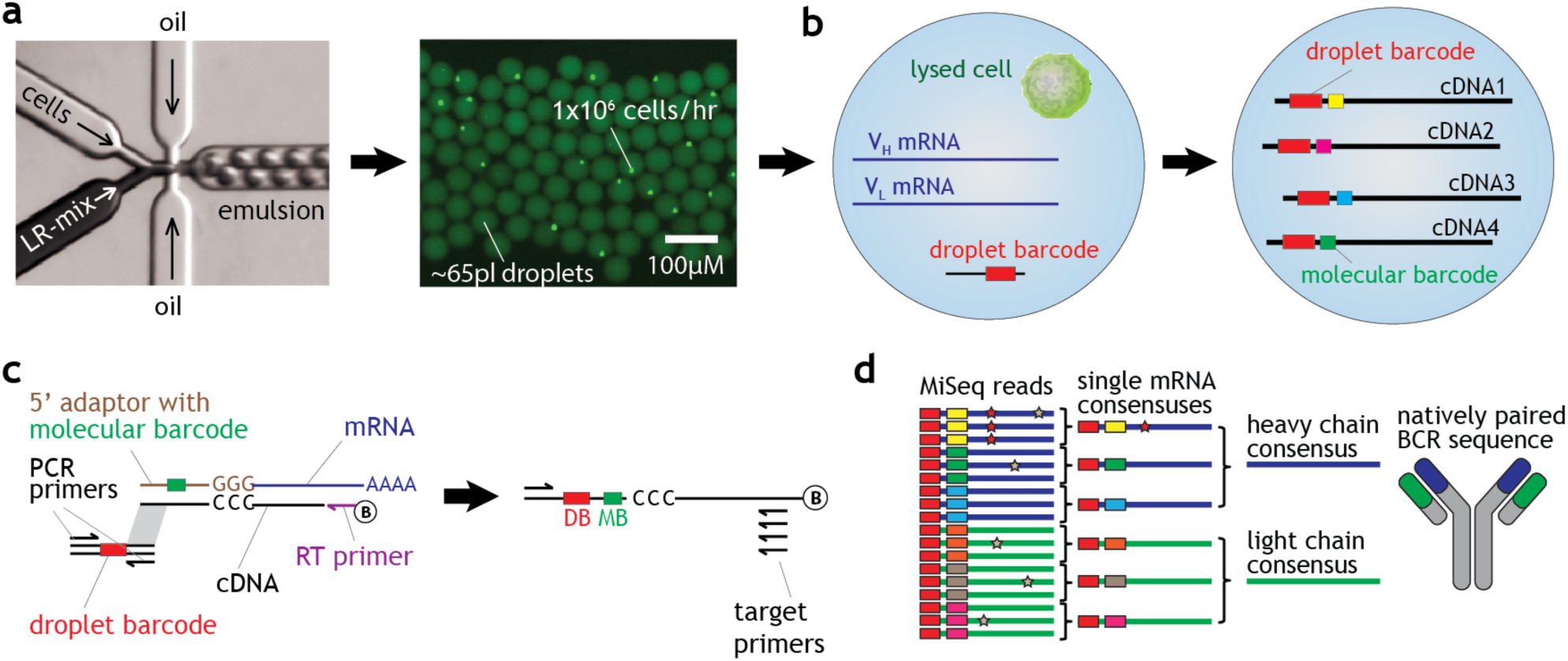
Single immune cell barcoding in emulsion. A) Two aqueous streams containing cells and lysis/reaction (LR) mix are passed into oil producing monodisperse emulsion at over 8 million droplets per hour. B) Within each droplet, cells are lysed and subjected to molecular- and droplet-specific barcoding in a single reaction. C) Target mRNA is first reverse transcribed and template switch-tagged with a universal adaptor sequence. Subsequently PCR amplification occurs of a droplet barcode template initially diluted to ∼1 molecule per droplet. Amplified barcodes are appended to template-switched cDNAs by complementary overlap extension. Products are recovered from the emulsion and purified using a biotin on the RT primer, before additional library processing steps and high throughput sequencing. C) Dual barcoding allows clustering of sequencing reads into their molecules and droplets of origin, reconstructing the native receptor chain pairings while minimizing sequencing errors and amplification biases.

## Large scale recovery of B-cell V_H_V_L_ pairs from a healthy blood sample

We initially developed the technology and assessed pairing capability and throughput with 3 million B-cells isolated by negative bead enrichment from peripheral blood of a healthy volunteer. The emulsion was split across six separate fractions which were processed in parallel and not remixed prior to sequencing. We loaded the emulsion at 0.2 cells per droplet, giving a Poisson expectation that ∼90% of occupied droplets contain single cells, which was consistent with emulsion droplet observations (Fig. 2a, Suppl. Fig. 2). After emulsion breaking and additional library processing steps we performed paired-end 325+300bp sequencing with Illumina MiSeq. To process the sequencing data, we used the droplet and molecular barcodes together to collect PCR replicate reads from each original mRNA molecule, and determined a consensus for each mRNA keeping only mRNA sequences built from at least two reads. We stitched forward and reverse reads to generate full-length products comprising the 5’ UTR, complete V(D)J sequence, and constant region sufficient for isotype determination. We then annotated rearranged immunoglobulin heavy and light chain sequences with IMGT High-VQuest^22^ and/or IgBLAST^23^.

**Figure 2.**
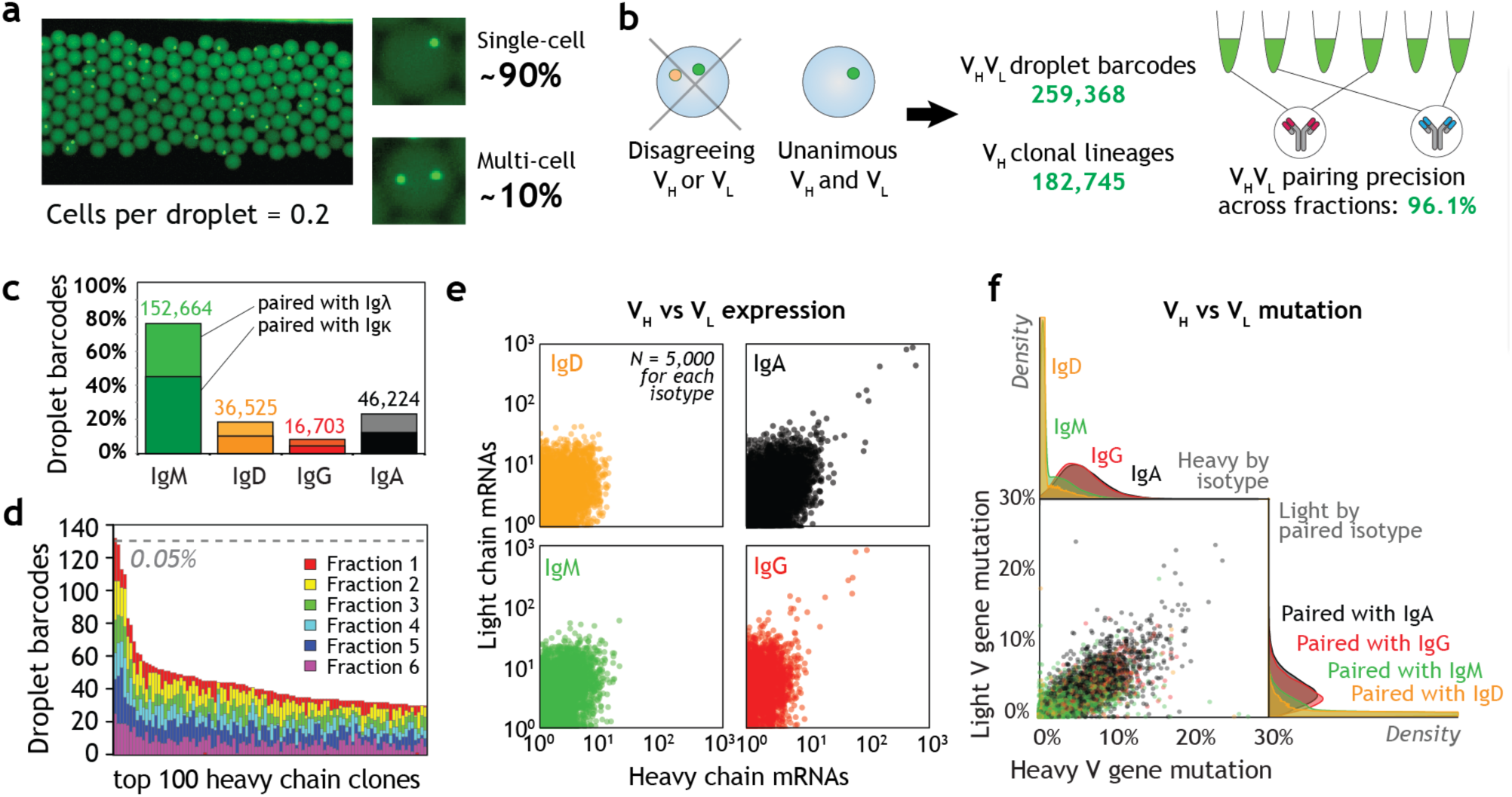
BCR recovery from isolated healthy B-cells. a) 3 million B-cells were passed into emulsion at 0.2 cells/droplet resulting in ∼90% of occupied cells containing single cells. b) After emulsion barcoding and sequencing, data was enriched for data from single-cell droplets and V_H_V_L_ pairing precision was estimated using pair consistency among expanded clones. c) Heavy chain isotype (most abundant isotype within each droplet) and light chain locus usage for 259,368 filtered V_H_V_L_ pairs. d) Rank abundance of the 100 most frequent heavy chain clones in each of six independent emulsion fractions. 0.05% overall frequency is marked. e) V_H_ vs V_L_ expression within cells as estimated by number of captured mRNAs within each droplet barcode. 5,000 points are shown for each isotype. V_H_ versus V_L_ mutation within pairs and distribution within each isotype.

The resulting dataset contained 324,988 droplet barcodes that were associated with at least one heavy chain (V_H_) and one light chain (V_L_) mRNA, with 229,869 distinct V_H_ clonal lineages present as estimated by heavy chain clustering analysis. Since this raw set includes data from multi-cell as well as single-cell droplets, we next enriched for data from single-cell droplets by filtering out droplet barcodes linked to non-unanimous heavy or light chain V(D)J sequences (Figure 2b). This step is made possible by the high diversity of typical immune repertoires, in which V_H_ or V_L_ mRNAs from two random cells will almost never match. The resulting enriched dataset comprised 259,368 V_H_V_L_ droplet barcodes and contained 182,745 V_H_ clonal lineages, representing comfortably the deepest sampling of a paired immune repertoire to date. We then directly estimated precision of the V_H_V_L_ pairings by identifying incidences of clonal expansion, since clonally related cells should show consistency in their V_H_V_L_ pairings^11^. We identified 2,604 V_H_ clones that were observed in more than one emulsion fraction with high confidence to be clonally expanded cells. The consistency of V_L_ sequences paired with these V_H_ clones across fractions indicated a pairing precision of 96.1% (see Online Methods), allowing high confidence in the entire filtered dataset of 259,368 V_H_V_L_ pairs. The cross-fraction V_H_ and V_L_ sequences were invariably associated with different droplet and molecular barcodes in each fraction and thus did not represent library cross-contamination. We also note that our analysis may underestimate pairing precision since some B-cells are known to express multiple light chains^11^.

75.0% of the 259,368 filtered V_H_V_L_ droplet barcodes or “V_H_V_L_ pairs” contained IgM and/or IgD (Figure 2c), which were frequently observed together as expected given the typical IgM^+^IgD^+^ phenotype of naïve B-cells. Lower but substantial fractions of IgA (18.3%) and IgG (6.6%) V_H_V_L_ pairs were also found. All V_H_ isotypes were paired with either Igĸ or Igλ in a ∼3:2 ratio. Among the 182,745 V_H_ clonal lineages we assessed clone expansion in two ways: the number of droplet barcodes associated with a clone, and observation of the clone across emulsion fractions. Clones seen in multiple droplet barcodes could reflect clonal expansion or multi-barcode droplets, which are expected in ∼37% of droplets given the initial λ = 1 Poisson dispersal of barcodes into droplets. However, any clone represented by > 8 droplet barcodes is likely to be genuinely expanded (Poisson probability in a single droplet < 10^-6^). While overall only 6.0% of clones were seen in more than one emulsion fraction, for the clones seen in more than 8 droplet barcodes (0.7% overall), 99% of them were seen in more than one fraction. The 100 most frequent clones (30-137 droplet barcodes each, Fig 2d) were all seen in at least five of six fractions. We conclude that a combination of barcode counting and independent fraction analysis allows detection of rare expanded lineages amongst a vast background of non-expanded clones^24^. Notably however, even the most abundant expanded clone was present at less than one cell in a thousand, exemplifying the huge diversity of human peripheral immune repertoires.

We next compared the number of captured mRNAs of each V_H_ and V_L_ chain within pairs as an estimate of expression level (Fig. 2e). While we generally captured less than ten heavy chain (mean 2.0) and light chain (mean 4.0) mRNAs per droplet barcode, a small population of droplet barcodes with dozens to hundreds of captured heavy and light chain mRNAs per cell was observed, almost exclusively from IgG and IgA expressing cells. Interestingly the degree of V_H_ and V_L_ mutation within pairs was strongly correlated both within each isotype (e.g. V_H_ vs V_L_ for IgG) and between isotypes (e.g. IgG vs IgM) (Fig. 2f). Furthermore, IgG and IgA pairs were almost all substantially mutated in both their V_H_ and V_L_ chains, whereas IgM and IgD pairs mostly showed little V_H_ or V_L_ mutation. These results are consistent with the mechanism of B-cell activation leading to class-switching from IgM and IgD to IgG or IgA, increased immunoglobulin expression and somatic hypermutation that affects both heavy and light chain loci in the cell. In addition to this observation that highly mutated V_H_ chains tend to be paired with highly mutated V_L_ chains, we conclude that our method is capable of generating large numbers of full-length, natively paired BCRs from resting human B-cell repertoires.

## Recovery of known low frequency V_H_V_L_ pairs from an HIV elite controller

As a further validation of the pairing sensitivity and accuracy of our assay, we processed a sample where several rare (< 1 cell in 10,000) native V_H_V_L_ pairings are already and publically known. We obtained peripheral B-cells from an HIV elite controller patient whose memory B cells have been mined heavily in recent years for antibodies displaying HIV neutralization activity^25,26^. 350,000 B-cells were processed to generate a total of 38,620 filtered V_H_V_L_ pairs. Interestingly, this individual showed a greater proportion of IgG than the previous healthy sample (Fig. 3a) or typical healthy peripheral B-cell repertoires^24,27,28^. We compared V_H_ sequences from this dataset to all reported broadly neutralizing antibodies (bNAbs) from this individual including PGT121^29^ and found eight close or identical V_H_ sequences, indicating that this family of bNAbs represents less than 0.03% of circulating B-cells. Crucially, all light chains paired to these heavy chains were of the expected and similarly rare bNAb lineage, displaying the same Igλ-V3-21/J3 rearrangement and hallmark triple codon insertion as previously reported^26^, supporting the high accuracy and sensitivity of our method. Furthermore, on a phylogenetic tree of all known and newly generated PGT121-like V_H_V_L_ pairs from this individual (Fig. 3b), the V_H_ and V_L_ trees show strikingly similar topology with paired V_H_ and V_L_ sequences occupying mirror-like positions, likely reflecting shared phylogenetic history as has been posited previously ^30^. The variant pairs discovered here fit well with this rule. Interestingly, two published antibodies PGT122 and PGT123 appear as exceptions; we did not find support for these two pairings but instead found PGT122V_H_:PGT123V_L_-like, and PGT123V_H_:PGT122V_L_-like pairs, addressing the unverified pairing in the original report. We synthesized DNA encoding the complete V(D)J regions of 8 novel PGT-like V_H_V_L_ pairs, expressed the antibodies as full IgG and tested their ability to neutralize multiple pseudostrains of HIV as previously described^26^ (Fig. 3c). The antibodies expressed well and all showed strong neutralizing activity against the virus, demonstrating the utility of our approach in rapidly generating natively paired functional antibody variants from a relevant biological sample.

**Figure 3.**
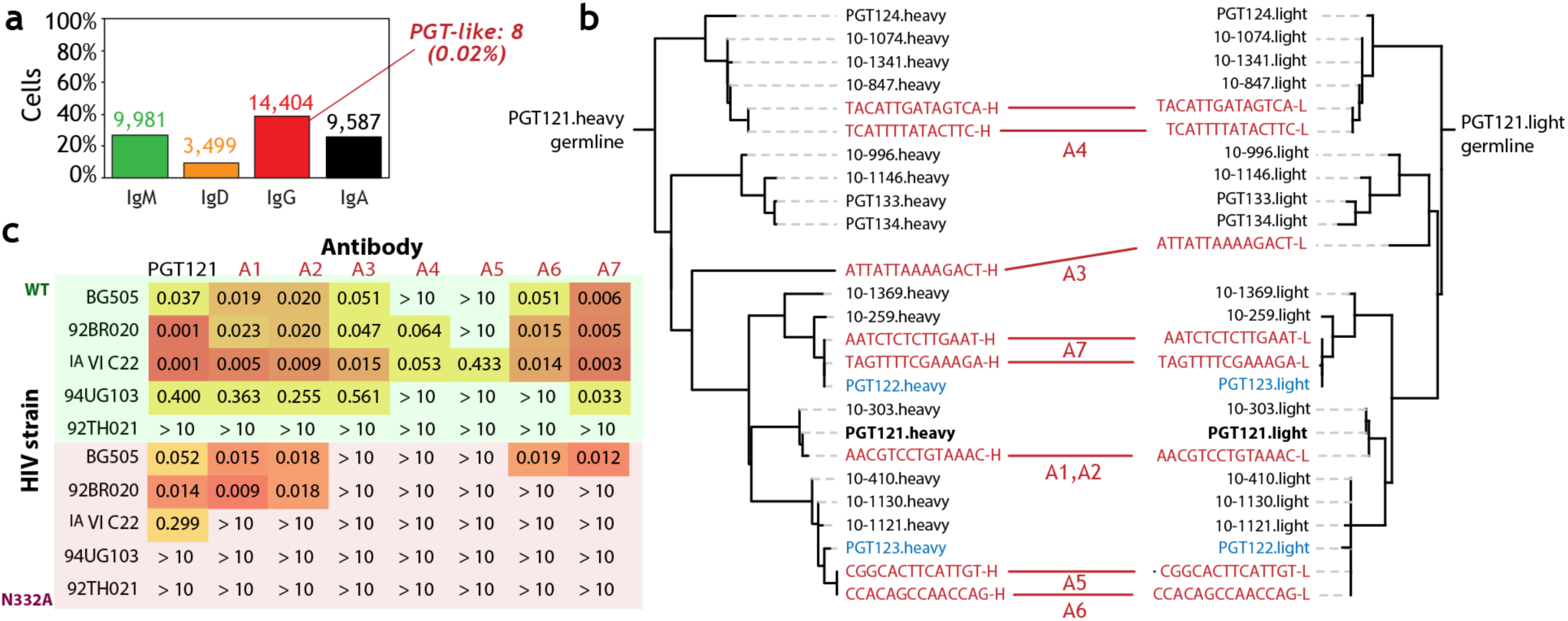
HIV bNAb discovery. B-cells from an HIV elite controller were entered into emulsion and BCR pairs were recovered. A) Heavy chain isotype distribution of the 38,620 recovered V_H_V_L_ pairs, where a rare proportion of the IgG chains aligned well to previously known bNAbs (“PGT-like”). B) Phylogenetic trees of complete VDJ amino acid sequences of known bNABs (black) plus the newly recovered ones (red, labeled with droplet barcode), with heavy (left) and light chains (right) plotted separately. Two antibodies potentially mispaired in current databases, PGT122 and PGT123, are blue. C) Neutralization activity (IC50, μg/mL) of the 8 newly discovered PGT-like variants against ten strains of HIV, compared to a control stock of PGT121.

## B-cell and T-cell receptor pairs from tumor infiltrating lymphocytes

Having validated our emulsion barcoding for high throughput recovery of paired receptors, we moved forwards with our primary objective of recovering immune receptors directly from a tumor. We took a protease-dissociated resected ovarian adenocarcinoma sample and entered 400,000 unsorted cells into emulsion. CD3/CD19 staining of a separate aliquot of the sample suggested substantial numbers of infiltrating B (∼5%) and T cells (∼20%) among the material. Single cell dispersal in the emulsion was similar to purified cells albeit with some limited clumping visible, and extensive variation in cell size and shape within the droplets as expected given the cell type heterogeneity of the sample (Fig. 4a). We included primers targeting the constant regions of T-cell receptor alpha and beta chains together with the BCR primers used previously, and following sequencing and stringent filtering recovered thousands of droplet barcodes linked to BCR or TCR products. To assess single cell precision we counted all possible combinations of the four target loci (V_H_, V_L_, V_α_, V_β_) within droplet barcodes (Fig. 4b). The vast majority (97.9%) of droplet barcodes with more than one target chain contained biologically expected pairings of BCR V_H_+V_L_ or TCR V_α_+V_β_ with only 2.1% containing mixed BCR-TCR combinations. Since barcoding of products is unbiased with respect to target chain, this result allows a high degree of confidence in the resulting 6,056 BCR V_H_V_L_ and 5,217 TCR V_α_V_β_ pairs. The BCRs showed striking dominance of IgG (>80%) compared to other isotypes (Fig. 4c), although all were present (IgE < 0.05% only). Kappa and lambda light chains were present in similar ratios to the peripheral blood datasets.

**Figure 4.**
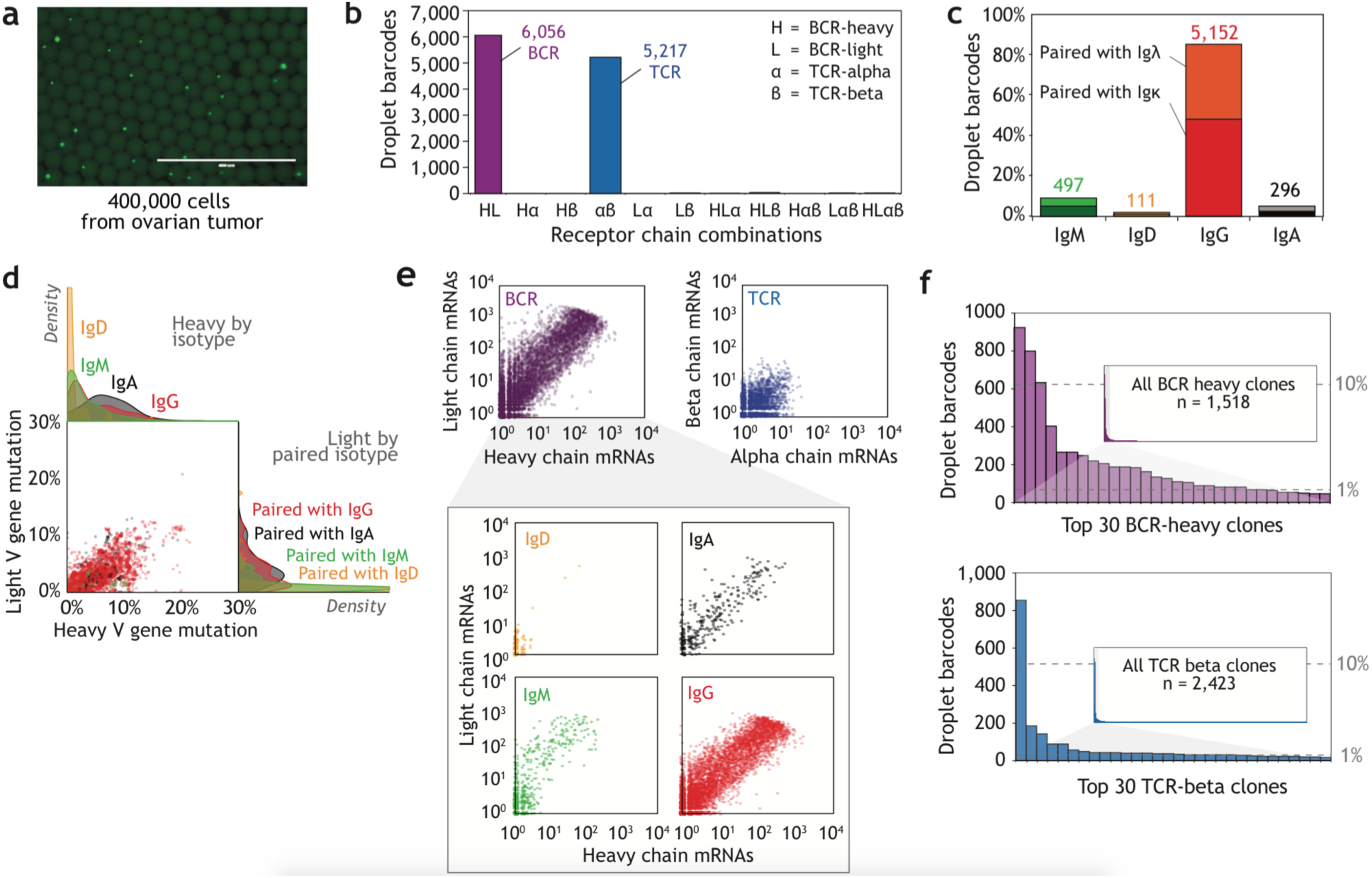
Characterization of TILs from an ovarian tumor. A) 400,000 unsorted dissociated cells from an ovarian tumor were entered into emulsion and BCR and TCR pairs were simultaneously recovered by emulsion barcoding. B) Numbers of all possible V_H_/V_L_/V_α_/V_β_ combinations observed within droplet barcodes after filtering. C) Heavy chain isotype distribution of the recovered BCR pairs. D) V_H_ vs V_L_ mutation correlation for BCR pairs and density distributions for each isotype. E) Numbers of captured mRNAs for TCR pairs and BCR pairs (overall and different isotypes). F) Clonal analysis and rank abundance plots for the recovered BCR and TCR pairs. 1% and 10% overall frequency levels are shown.

Similarly to peripheral blood we observed a correlation between BCR isotype and mutation level of both V_H_ and V_L_ chains, with IgG and IgA pairs showing greater V_H_ and V_L_ mutation than IgD and IgM, and a general correlation of mutation between V_H_ and V_L_ within each isotype (Fig 4d). Interestingly, while IgD, IgM and IgA pairs showed very similar mutational distributions between the tumor and peripheral blood datasets (Fig 2f), the tumor IgG fraction also contained a substantial proportion of little-to non-mutated sequences that we did not observe in the peripheral blood. For TCRs, and for BCRs containing IgD, we observed similar numbers of captured mRNAs per droplet barcode to the BCR results from peripheral blood (Fig. 4e, mostly <10 per droplet barcode). In stark contrast, the tumor-derived IgM, IgA and IgG pairs showed a 10 to 100-fold increased average expression level with hundreds or thousands of target mRNAs captured in many of the droplet barcodes. We then assessed the diversity of the captured TIL-TCR and BCR repertoires (Fig. 4f). Among the 5,217 total TCR pairs we observed 2,423 distinct TCR beta clones. Seven clones were present at a frequency >1% with the top clone representing 16.9% of all droplet barcodes. Among the 6,056 total BCR pairs we observed 1,518 distinct heavy chain clones, with 15 clones at >1% frequency but none >5%. While this represents substantially more restricted diversity than the healthy peripheral BCR repertoire (where no clone was present in greater than 0.06% frequency), the presence of so many class-switched, mutated and highly expressed clones in the tumor sample demonstrates the necessity of a deep and sensitive sampling approach for TIL characterization. We conclude that our method allows rapid retrieval of large numbers of TIL immune receptor pairs, from both B and T cells simultaneously, without the need for prior sorting or exogenous activation of defined TIL populations.

## Capture of additional phenotypic markers of interest

Pairing of receptor chains by droplet barcoding potentially allows capture of additional targets besides immune receptors. To investigate this possibility we separated healthy T-cells into CD4^+^ and CD8^+^ populations by magnetic bead enrichment and entered 20,000 cells of each type into separate emulsion runs, with primers targeting TCR alpha and beta chains and CD4 and CD8 mRNAs. After sequencing, 47.0% of 3,861 droplet barcodes containing TCR V_α_ and V_β_ (“TCR pairs”) from CD4^+^ isolated cells were linked to CD4 mRNA, while only 0.3% were linked to CD8 mRNA. Conversely, 50.6% of 2,235 TCR pairs from CD8^+^ isolated cells were associated with CD8 mRNA, while only 0.6% were linked to CD4 mRNA (suppl. Fig 3). This demonstrates the high specificity but limited sensitivity of an mRNA-based approach to cell phenotyping, similar to a previous report^31^. In contrast, proteins such as cell surface receptors are usually present in far higher numbers (1,000-100,000 per cell) than their coding mRNAs, potentially making them easier to detect as well as being potentially more directly relevant to cell phenotype. To measure target protein levels on each cell we conjugated custom oligonucleotide DNA labels to anti-human CD4 and CD8 antibodies, and incubated the labeled antibodies with an unseparated mixture of CD4^+^ and CD8^+^ T-cells before entry of 30,000 cells into an emulsion (Fig. 5a). The DNA labels carry antibody-specific sequence tags as well as molecular barcodes and sequence complementarity to the amplified droplet barcodes, allowing emulsion droplet barcoding and molecular counting similarly to that done for mRNAs. We targeted the DNA labels as well as TCR, CD4 and CD8 mRNAs simultaneously. After sequencing and filtering we identified 3,682 droplet barcodes with high confidence TCR V_α_V_β_ pairs. Consistent with the previous experiment, roughly half (52%) of the TCR pairs could be assigned CD4 or CD8 status based on mRNA (Figure 5b). However, over 95% of droplet barcodes could be assigned CD4 or CD8 based on protein status, with average molecular counts per droplet considerably higher for CD4/8 proteins (mean 20.5) than CD4/8 mRNAs (mean 1.0). Concordance between mRNA and protein signals was high (Fig 5.c): 96.0% of droplets given both mRNA and protein calls were in agreement. In some rare instances, both CD4 and CD8 proteins were detected, possibly a result of droplets that contained two or more cells. Adding to recent studies reporting similar approaches ^32,33^, we conclude that emulsion barcoding allows direct linking of single cell mRNA and protein markers of interest, here with the added component of full-length immune receptor sequences, all at high throughput. Application of this approach to TILs with an expanded, immune-oncology relevant marker set such as anti-PD-1 and anti-CTLA-4 is now warranted.

**Figure 5.**
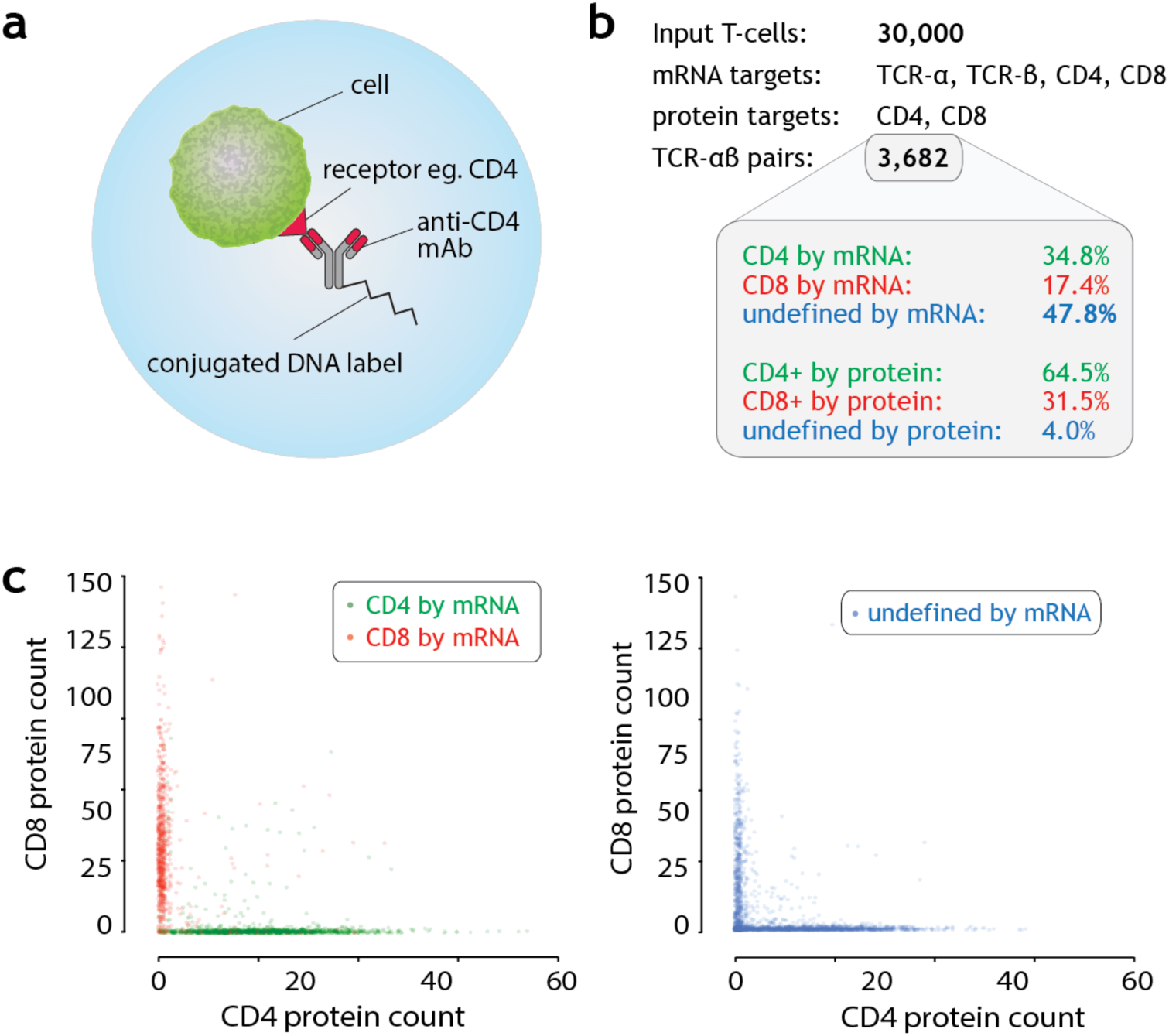
Co-capture of immune receptor sequences with additional mRNA and protein targets. A) Surface protein targets are quantified by pre-incubating cells with DNA-labeled staining antibodies prior to emulsion sequencing. B) CD4 and CD8 mRNA and protein detection in 3,682 droplet barcode TCR V_α_V_β_ pairs generated from healthy human T-cells. Undefined = could not be called CD4 or CD8 by molecular count majority rule. C) Concordance between mRNA and protein measurements. Each point is a droplet barcode linked to a TCR V_α_V_β_ pair, colored according to its CD4 or CD8 assignment by mRNA.

## Discussion

In this study we processed several million unstimulated naïve and resting memory B-cells from peripheral blood and hundreds of thousands of dissociated tumor cells, yielding repertoire-level datasets of full-length, paired BCRs from the blood sample and both BCRs and TCRs from TILs in the tumor. While T-cells have traditionally been the primary focus of TIL research, the roles of B-cells in tumors are receiving increasing attention^34,35^. Renewed interest in TILs from the broad scientific community can be attributed to groundbreaking development in cell transfer immunotherapies^36^, development of CAR-T against solid tumors^37^, and the discovery of checkpoint inhibitors and their impact on tumor growth^8^. These fields of research all require an understanding of the TILs involved, either to monitor TIL fluctuation and evolution in responding versus non-responding patients, or for the discovery of novel tumor antigens using TIL receptors as a tool. A major barrier to advancing this field of research is that only a small percentage of TILs in a tumor may be antigen specific, with a large background of non-specific immune cells present due to general inflammation^38,39^. Therefore it is critical to sample sufficient numbers of TCRs and BCRs to identify those specifically enriched in the tumor environment, a task likely to require many thousands of cells rather than the small numbers afforded by plate-based TIL analysis^31^. In addition, since TILs have been reported to show non-canonical surface phenotypes^40^, marker-based sorting may miss important TIL populations. The task of single cell sequencing from unsorted tumor tissue therefore requires an extremely high throughput single cell approach. While several recent studies have reported compelling progress in scale-up of single cell transcriptome analysis, using barcoded oligonucleotide beads in emulsion droplets^16–19^, to date these have not demonstrated the read length^19^ or throughput of several hundred thousand cells per sample^16–18^ necessary for meaningful V(D)J capture of immune repertoires from heterogeneous samples. Our method in contrast affords higher throughput through the use of emulsion droplets 15-75 times smaller than used previously, a size previously inaccessible due to highly inhibitory RT-PCR conditions, and the generation of clonal droplet barcodes within the emulsion by PCR, thus removing the need for prior synthesis of barcoded beads. In addition, since the method presented here involves both reverse transcription and PCR in the original emulsion droplets, a greater range of target types can be aimed at allowing mRNA, labeled protein and potentially genomic DNA to be captured simultaneously from single cells, while also minimizing the risk of post-emulsion cross-contamination between barcodes^17^.

To date, all methods for targeted BCR or TCR sequencing suffer from at least one of the following limitations: unpaired receptor chains^21^, low throughput of paired chains (<10^4^ per experiment^41,42^), or partial sequence recovery^43^. The technical challenge is particularly high since primary uncultured immune cells can contain 100-fold less receptor RNA than *in vitro* stimulated cells^44,45^. In contrast the emulsion method presented here overcomes these limitations and displays a number of additional advantages. Firstly, linking the chains with short sequence barcodes, rather than physically splicing into long amplicons as performed by Dekosky et al.^11^ allows sequencing of the complete V(D)J variable region, ensuring comprehensive analysis of somatic mutations and allowing fully accurate functional reconstruction. Secondly, by dual barcoding both the mRNAs themselves and all products from a single cell, PCR and sequencing errors can be minimized and in particular the number of cells in each clonal lineage can be deconvoluted from the expression level within cells of that clone. This feature, applied here for the first time to immune repertoire analysis, overcomes the limitations of immune sequencing from bulk mRNA, which measures overall expression but not clonal abundance, as well as from genomic DNA, which reveals clonal abundance but nothing about expression^46^. Third, our barcoding approach can be extended to additional genes of phenotypic value at both the mRNA and protein level, adding important information about the sub-types of T- and B-cells expressing each receptor sequence. To our knowledge we present here the first demonstration of simultaneous immune receptor sequencing with protein measurements from single cells in emulsion. This approach combines the type of information collected by flow cytometry with the digital resolution of immune sequencing, allowing direct linking of each cell’s phenotype to its full-length receptor sequence. The method therefore promises to be a powerful addition to tumor receptor profiling, by revealing which T- and B-cell lineages represent particular cell subtypes and states such as immune suppression or “exhaustion”, often a key indicator of anti-tumor reactivity^8^. The approach could also be applied to discovery of antibodies specific for a desired antigen, by incubating B-cells with a DNA-barcoded target antigen prior to emulsion.

In summary, in this study we recovered natively paired BCRs and TCRs from thousands of TILs in a dissociated tumor sample, leading to identification of dozens of clonally expanded receptor lineages that are mutated, class-switched and highly expressed by the cells, strongly suggesting the specific recruitment of functionally active B and T cells to the tumor site. Taken alongside complementary approaches^47,48^ to identify the targets of TIL receptors found in this and other tumors, the technology described here should enable the discovery of novel tumor antigens, further our basic understanding of tumor immuno-biology, and provide a powerful tool to investigate the immune effects of emerging immunotherapies.

## Acknowledgements

The authors wish to thank Uri Laserson, Teresa J. Broering and David Fabrizio for technical discussions.

## Author Contributions

A.W.B., S.J.G. and F.V. designed and developed the emulsion sequencing method with contributions from C.R.C, B.J.B. and M.V.T.; D.S. performed neutralization experiments; S.T. designed and directed the data analysis; S.T., A.W.B., S.J.G., D.K., J.V.H. and B.J.B., analyzed data; D.R.B. and G.M.C., provided technical expertise; F.V. supervised the study; A.W.B., S.T., S.J.G. and F.V. wrote the manuscript with input from all authors.

## Supplemental Information

Supplemental information representing 3 supplementary figures, 1 supplementary table and supplementary methods is provided in a separate file.

## Human Samples

The blood sample for healthy repertoire validation was collected under the approval of the Personal Genome Project^49^. PBMCs for the HIV bNAb experiment were obtained from donor 17, an HIV-1 infected donor from the IAVI Protocol G cohort ^50^. All human HIV samples were collected with written informed consent under clinical protocols approved by the Republic of Rwanda National Ethics Committee, the Emory University Institutional Review Board, the University of Zambia Research Ethics Committee, the Charing Cross Research Ethics Committee, the UVRI Science and Ethics Committee, the University of New South Wales Research Ethics Committee. St. Vincent’s Hospital and Eastern Sydney Area Health Service, Kenyatta National Hospital Ethics and Research Committee, University of Cape Town Research Ethics Committee, the International Institutional Review Board, the Mahidol University Ethics Committee, the Walter Reed Army Institute of Research (WRAIR) Institutional Review Board, and the Ivory Coast Comite “National d’Ethique des Sciences de la Vie et de la Sante” (CNESVS). Cryopreserved, dissociated resected ovarian adenocarcinoma from a single donor was obtained from Conversant Biologics with written informed consent under an IRB approved protocol.

## Methods

### Cell preparation

For the study of 3 million healthy B-cells, 50 ml blood was drawn into Vacutainer CPT Cell Preparation Tubes with sodium heparin (BD), centrifuged for 20 min at 1800 × *g*, washed twice in cell preparation buffer (1x PBS supplemented with 2% fetal bovine serum and 2mM EDTA), using spins at 200 × *g* to remove platelets, and the resulting PBMCs were cryopreserved in RPMI-1640 medium (Life Technologies) + 20% fetal bovine serum + 10% DMSO at -80 °C until needed. Prior to emulsion generation, PBMCs were thawed, washed twice in cell preparation buffer and counted. B-cells were isolated using a negative selection-based human B-cell enrichment kit (Stem Cell Technologies) according to the manufacturer’s instructions. Cells were passed through a 20 μm cell strainer and diluted to 6.2 x 10^6^ cells/ml (3-million B-cell experiment) or 3.1 x 10^6^ cells/ml (PGT-donor and ovarian tumor experiments) in cell preparation buffer.

### Immune receptor barcoding in emulsion

The emulsion generation platform consisted of three Mitos P-Pumps (Dolomite Microfluidics) driven by a single air compressor, each with a Mitos Flow Rate sensor, to allow computer-controlled flow of two aqueous phases and one fluorophilic oil continuous phase into a fluorophilically-coated quartz Dolomite Small 2-Reagent chip. One aqueous input channel contained the cells at the required density to produce the desired cells-per-droplet occupancy level, while the second aqueous channel contained lysis and reaction mix, consisting of AbPair reaction buffer and oligonucleotides (suppl. Table 1,), 5 units/μl MuMLV-based reverse transcriptase (Thermo Scientific) and 0.1 units/μl Herculase II PCR polymerase (Figure 1). A 100-μL Hamilton Microliter syringe was used to overload a 100-μL internal diameter PEEK tubing sample loop in two injections of ∼100 μL each of LR mix. A 100-μL Hamilton Gastight syringe was used to load ∼110 μL of the cell suspension into a ∼100-μL, 0.2-mm internal diameter FEP tubing loop. The emulsion was formed by focused flow jetting of the aqueous phases at identical flow rates through the 2-reagent chip with simultaneous oil flow from the two oil channels in the chip. The emulsion leaving the chip exit channel was dripped into 0.2-ml PCR strip tubes (Eppendorf) on a cold block, after which excess oil was removed by pipetting from the bottom of the tube, 40ul of overlay solution was added (25 mM Na-EDTA, pH 8.0) and tubes were transferred to a standard thermocycler for the transcript tagging reaction. During a 45-min reverse transcription (RT) step, RNA is reverse transcribed at 42 °C with target-specific RT primers (Supplementary Table 1), with template-switch-based addition of a universal adaptor sequence containing a randomized molecular barcode as previously described^21,51^. Following RT, emulsions were subjected to 40 cycles of thermocycling (each cycle: 82C for 10 sec, 65C for 25 sec) to perform PCR amplification of the droplet barcode templates, which were diluted in the initial lysis and reaction mix to 30,000 cp/μl, generating a concentration in the final mixture of 15,000 cp/ul or ∼1 per ∼65pl droplet. One end of the droplet barcode comprises the Illumina read 2 (“P7”) primer site, whereas the other end matches the common sequence of the universal adaptor oligonucleotide. Therefore, during PCR, template-switched cDNAs can anneal to amplified droplet barcode strands and become spliced by overlap extension to produce full-length products containing target, molecular barcode and droplet barcode sequences.

### Emulsion breaking, cleanup, downstream PCRs, pooling and sequencing

After thermocycling, the overlay solution was removed by pipetting and 40ul emulsion breaking solution (1:1 FC-40:perfluorooctanol) were added together with 15ul lysate clearing solution (12.5 μL Qiagen Protease, 2.5 μL 0.5 M Na-EDTA, pH 8.0). After inverting 10 times to break the emulsion, the mixture was incubated for 15 minutes at 50 °C and 3 minutes at 95 °C to inactivate the protease. After centrifugation at 15,000 × *g* for 1 min to isolate the aqueous phase, the recovered material was rigorously purified to remove oligonucleotides, reagents and excess droplet barcode PCR products. Since full length products contain biotin due to 5’ biotinylation of the RT primer, they can be efficiently separated from excess droplet barcode PCR products by cleanup on streptavidin beads, thus minimizing downstream PCR recombination artifacts, a common problem in extension-by-overlap approaches^52^. First products were purified using AMPure XP beads (Agencourt) using manufacturer’s instructions at a 1:1 ratio, followed by cleanup using streptavidin beads (New England Biolabs) also using manufacturer’s instructions, followed by elution in deionized water at 95 °C^53^, followed by a second cleanup with AMPure XP beads at a 1:1 ratio. Products were then entered into a target enrichment PCR in which primers specific to the constant regions of the B- or T-cell receptor targets (Supplementary Table 1) were used together with a primer specific to the universal end of the droplet barcode sequence. This reverse primer also contained a six-base index barcode for multiplexed sequencing on the MiSeq instrument according the manufacturer’s instructions. Thus, only full-length, droplet-barcoded target sequences are amplified in this step. We first amplified all targets together for seven cycles of 98C 10 seconds; 64C 20 seconds; 72C 15 seconds, using Q5 Hot Start polymerase (New England Biolabs) under manufacturer-recommended conditions, including a 2 minute 98C polymerase activation step at the beginning of the reaction. This was followed by AMPure XP cleanup at a 1.5:1 beads:PCR ratio. We then performed a second seven-cycle targeting each chain (VH, VL, VA, VB) separately, using the same thermocycling conditions as before, followed by AMPure XP cleanup. We then performed a final PCR with the same thermocycling conditions and 5–15 cycles (depending on yield as judged by qPCR) to add the full-length Illumina sequencing adaptors and generate enough material for TapeStation D1000 (Agilent) quantification. We then pooled libraries and sequenced on the V3 2 x 300bp MiSeq platform (Illumina).

### Modifications to the standard MiSeq platform

Reconstruction of the complete variable V(D)J region of BCR or TCR requires stitching the two paired-end Illumina reads. To improve this process we extended the forward read of the 2 x 300bp kit to 325bp. We used 10% phiX spike-in to alleviate issues of limited library diversity, since immune receptor libraries have limited diversity in the constant region primer sites.

### Overview of bioinformatic processing of reads

Illumina MiSeq reads were processed using custom pipelines built around the pRESTO package (version 0.4) ^54^ to generate full length consensus sequences for mRNA molecules from each droplet, annotated with IgBLAST and/or IMGT/HighV-QUEST, and further aligned, filtered, and processed with custom scripts and the Change-O package ^55^ to generate statistics. Detailed methods can be found in Supplementary Material.

### Read processing and annotation, Isotype assignment

Raw read processing, V(D)J annotation and clonal assignment was performed with custom pipelines utilizing the pRESTO and Change-O packages. Briefly, raw Illumina paired-end 325+300 bp reads were quality-controlled, primer-trimmed, and droplet-specific (DB) and molecule-specific barcodes (MB) identified via fuzzy matching of primer sites. Together, DB and MB uniquely specify a molecule of origin, and this unique molecular identifier (UMI) was used to group agreeing PCR replicate reads (minimum of two) hailing from the same molecule to generate a consensus for each mRNA sequence. Isotype-specific priming was confirmed by fuzzy matching of known isotype-specific constant regions within primer-trimmed sequences. V(D)J germline segments and rearrangement structure was determined using IgBLAST and confirmed with IMGT/HighV-QUEST^22^ where appropriate, parsed by Change-O and custom scripts. Clones were assigned via single-linkage clustering within groups of functional V(D)J sequences having matching IGHV gene, IGHJ gene, and junction length as implemented in Change-O. For the 3 million circulating cell dataset, a weighted intraclonal distance of 4.0, using a symmetrized transition/transversion model^56^ as previously described^57^ was used as the nearest-neighbor distance cutoff within clones (see Supplementary Material).

### Droplet immune receptor inclusion filtering and pairing fidelity calculation

Precision of B-cell sequence recovery from droplets can be assessed in two ways with this barcoding method: using intra-droplet mRNA sequence agreement, and via cross-fraction pairing agreement. Within each droplet, multiple mRNAs are captured per locus; expressed V(D)J sequences from one cell should agree. The presence of more than one productive VDJ and one productive VJ sequence per droplet is flagged bioinformatically as putative immune receptor inclusion or multi-cell occupancy, using a cutoff of 2% sequence diversity (mean pairwise nucleotide differences pi^58^ <0.02) of multiple aligned V(D)J segments to define sequence agreement. Heavy and light chain consensus sequences were built for each allelically excluded droplet, and were used for clone definition and cross-fraction pairing analysis. For the 3 million circulating B-cell dataset, each V_H_ lineage is associated with one (in the ideal case) of >20,000 light chain clones in the dataset. Among 259,368 immune-locus-excluded droplets with VHVL pairs, 10,870 VDJ heavy locus rearrangement clusters were present in at least two of six physically separated emulsion fractions. These clusters represent either expanded lineages or independent but similar rearrangement of the same VJ exons. Where a VDJ rearrangement is paired with a consistent VJ rearrangement across two replicates, both experiments independently produced a true positive (33,157 of 35,922 possible pairwise comparisons for 2,604 clones with rarer rearrangements; see Supplementary Material). Thus, the precision for each replicate is 96.1% (0.923^0.5).

### HIV bNAb candidate sequence discovery

New natively paired broadly-neutralizing antibodies (BNAbs) to HIV were discovered by mining our 38,620 V_H_V_L_ pairs for similarity to known bNAb HCDR3s, VDJ sequences and Donor 17 lineages culled from the literature^29^ using tblastx^59^, MUSCLE^60^, and PhyML^61^, followed by manual inspection of phylogenetic trees of full V(D)J amino acid sequence to select antibody candidates interspersing with known bNAb sequences.

### HIV bNAb protein expression and purification

Antibody sequences were synthesized and cloned into previously described heavy and light chain vectors^62,63^. Heavy and light chain plasmids were co-transfected (1:1 ratio) in 293 FreeStyle cells using 293fectin (Invitrogen) according to the manufacturer’s protocol. Antibody supernatants were harvested four days following transfection and purified by protein A affinity chromatography. Purified antibodies were buffer exchanged into PBS before use in further assays.

### Pseudovirus production and neutralization assays

Pseudoviruses were generated by transfection of 293T cells with an HIV-1 Env expressing plasmid and an Env-deficient genomic backbone plasmid (pSG3ΔEnv), as described previously^64^. Pseudoviruses were harvested 72 hr post-transfection for use in neutralization assays. Neutralizing activity was assessed using a single round of replication pseudovirus assay and TZM-Bl target cells, as described previously ^64^. Briefly, TZM-bl cells were seeded in a 96-well flat bottom plate. To this plate was added pseudovirus, which was preincubated with serial dilutions of antibody for 1 hr at 37 °C. Luciferase reporter gene expression was quantified 72 hr after infection upon lysis and addition of Bright-Glo Luciferase substrate (Promega). To determine IC50 values, dose-response curves were fit by nonlinear regression.

### Ovarian tumor target chain identification

After simultaneous BCR and TCR capture from ovarian dissociated tumor tissue in emulsion, we filtered reads using molecular and droplet barcodes as previously described, but then looked for the presence of each of the four possible target chain types (BCR V_H_, BCR V_L_, TCR V_α_, TCR V_**β**_). Target chains were retained if they were supported by at least two mRNAs, each with at least two sequencing reads. All droplet barcodes containing only BCR V_H_V_L_ or TCR V_α_V_β_ pairs were analyzed further. BCR heavy chain and TCR beta chain clones were called based on distinct CDR3 amino acid sequences.

### Protein detection in emulsion through DNA-labelled antibody staining

Single-stranded, 200 bp DNA oligonucleotides were designed to contain unique 5 bp antigen ID sequences and were modified with a 5’ amino group (Integrated DNA Technologies). Mouse monoclonal, anti-human CD4 (BioLegend, 300516) and CD8a (BioLegend, #301018) antibodies were conjugated to DNA oligonucleotide tags using the Thunder-Link kit (Innova Biosciences) according to manufacturer’s protocol. For cell labeling prior to emulsion, two million negatively selected T-cells from peripheral blood were diluted in 400 uL cell buffer + 2 mM EDTA + 0.05% sodium azide. Single stranded salmon sperm DNA was added to cells to a final concentration of 200 ug/mL and cells were rotated at room temperature for five minutes. A mixture of CD4 and CD8a DNA-labeled antibodies (each to a final concentration of 5 nM) was added to the cells and incubated at room temperature for 30 minutes. Cells were washed three times with cell Buffer + 2 mM EDTA + 0.05% sodium azide + 200 ug/mL single stranded salmon sperm DNA. Cells were resuspended in cell buffer + 0.05% sodium azide prior to entry into emulsion analysis. 30,000 cells were used for emulsion sequencing.

